# Loss of *Cochlin* drives impairments in tendon structure and function

**DOI:** 10.1101/2024.11.14.623674

**Authors:** Emmanuela Adjei-Sowah, Elsa Lecaj, Neeta Adhikari, Clara Sensini, Anne E.C. Nichols, Mark R. Buckley, Alayna E. Loiselle

## Abstract

Aging tendons undergo disruptions in homeostasis, increased susceptibility to injury, and reduced capacity for healing. Exploring the mechanisms behind this disruption in homeostasis is essential for developing therapeutics aimed at maintaining tendon health through the lifespan. We have previously identified that the extracellular matrix protein, *Cochlin*, which is highly expressed in healthy flexor tendon, is consistently lost during both natural aging and upon depletion of Scleraxis-lineage cells in young animals, which recapitulates many aging-associated homeostatic disruptions. Therefore, we hypothesized that loss of Cochlin would disrupt tendon homeostasis, including alterations in collagen fibril organization, and impaired tendon mechanics. By 3-months of age, *Cochlin^-/-^* flexor tendons exhibited altered collagen structure, with these changes persisting through at least 9-months. In addition, Cochlin*^-/-^* tendons demonstrated significant declines in structural and material properties at 6-months, and structural properties at 9-months. While *Cochlin^-/-^* did not drastically change the overall tendon proteome, consistent decreases in proteins associated with RNA metabolism, extracellular matrix production and the cytoskeleton were observed in *Cochlin*^-/-^. Interestingly, homeostatic disruption via *Cochlin^-/-^* did not impair the tendon healing process. Taken together, these data define a critical role for *Cochlin* in maintaining tendon homeostasis and suggest retention or restoration of *Cochlin* as a potential therapeutic approach to retain tendon structure and function through the lifespan.

## Introduction

Tendon homeostasis is a finely tuned process that maintains the structural and functional integrity of the tendon over time ^1–5^. Disruptions to this homeostatic balance can result in alterations to the tendon extracellular matrix (ECM), including changes in ECM composition, organization, and material quality, which impair the tendon’s ability to respond to mechanical loading and repair after injury ^2–5^. These disruptions are often linked to tendon pathologies, such as tendinopathy and ruptures, which hinder overall musculoskeletal function^6–8^. The maintenance of tendon homeostasis relies on the balance between ECM synthesis and degradation, a process regulated largely by resident tenocytes that respond to mechanical and biochemical cues ^9^. We have previously shown that during natural aging, tendon homeostasis is disrupted due in part to loss of tenocytes with strong biosynthetic functions^4^. Moreover, our previous work demonstrated that upon depletion of Scleraxis-lineage (Scx^Lin^) cells in 3-month-old mice, homeostatic disruptions similar to those observed with natural aging are observed, including increased collagen disorganization ^2, 4^. Comparative proteomic analysis between young, aged (21 months) and Scx^Lin^-depleted tendons identified consistent loss of multiple high turnover rate ECM proteins during disrupted tissue homeostasis. Among these, Cochlin, a non-collagenous ECM protein, emerged as a potential regulator of tissue homeostasis ^4^.

*Cochlin* is the primary non-collagenous ECM protein found in the inner ear ^10–12^. Although its precise function is not completely understood, it is believed that mutations in the *COCH* gene cause the accumulation of misfolded protein, leading to progressive, late-onset hearing loss ^13–16^, with *Cochlin*^-/-^ mice demonstrating this phenotype^17^. Moreover, *Cochlin* is expressed in the spiral ligament of the inner ear, and loss of *Cochlin* resulted in degeneration of the spiral ligament ^18^. Despite its recognition as an important ECM protein in other tissues, the role of *Cochlin* in tendon remains largely unknown.

In this study, our primary goal was to identify the structural and compositional changes resulting from loss of *Cochlin* in young adult flexor tendons. We hypothesized that loss of *Cochlin* would drive dysregulated ECM homeostasis, leading to compromised tendon function. By assessing the structural, mechanical, and molecular changes that occur with loss of *Cochlin*, we have defined the requirement for *Cochlin* in maintaining tendon homeostasis, identifying Cochlin retention as a potential means to retain tendon structure-function through the lifespan.

## Materials and methods

### Animal ethics

All studies were carried out in strict accordance with the recommendations in the Guide for the Care and Use of Laboratory Animals of the National Institutes of Health. All animal procedures were approved by the University Committee on Animal Research (UCAR) at the University of Rochester (# 2014-004E).

### Mice

B6.129S1(Cg)-Cochtm1.1Stw/YuanJ (#024691) were purchased from Jackson laboratory and were bred to generate *Cochlin^-/-^* and wildtype (WT) littermates. Mice were used at 3, 6, and 9 months of age. Equal distributions of male and female mice were used for all studies.

### Transmission Electron Microscopy (TEM) Imaging and Analysis

To isolate FDL tendons for TEM, the hind paw was fully submerged in TEM fixative (2.5% glutaraldehyde/4.0% paraformaldehyde buffered in 0.1 M sodium cacodylate), and tendons (N = 6 per age per genotype) were carefully isolated to ensure preservation of tendon ultrastructure and further fixed in TEM fixative, after which they were post-fixed in buffered 1% osmium tetroxide/1.5% potassium ferrocyanide, dehydrated in a graded series of ethanol to 100%, transitioned into ethanol/propylene oxide (1:1), propylene oxide only, then propylene oxide/Eponate/Araldite epoxy resin (2:1 then 1:1), and finally 100% epoxy resin overnight in the Electron Microscopy Shared Resource Laboratory. The next day the tendons were placed into silicone molds with fresh epoxy resin and polymerized for 48 hours at 60°C. The epoxy blocks were trimmed for axial one-micron sections that were stained with Toluidine blue. An ultramicrotome with a diamond knife was used to cut 70 nm ultrathin sections for placement onto formvar/carbon coated copper slots grids then stained with uranyl acetate and lead citrate. The grids were imaged using a Hitachi 7650 TEM and an AMT 12-megapixel digital camera. Three distinct regions were imaged from the mid-substance of each tendon, at 40,000x magnification. One image per sample was used for measurement of fibril density and diameter; a straight line was drawn along the x-axis of each fibril within each image in ImageJ software. Only fibrils exhibiting a complete circumference were included in the measurements, and density was calculated as fibrils per µm^2^.

### Sample Preparation for Mass Spectrometry

Tendons were identified and carefully isolated to ensure that no connective tissue or muscle was collected, after which it was immediately flash frozen in liquid nitrogen and transported to the Mass Spectrometry Resource Lab. Each tendon was homogenized in 100 µL of 5% SDS, 100 mM TEAB, vortexed and subsequently sonicated (using QSonica) for 5 cycles, with a 1-minute resting interval on ice following each cycle. The lysates were centrifuged at 15,000 x g for 5 minutes to gather cellular debris, and the resulting supernatant was collected. A BCA assay (Thermo Scientific) was used to determine protein concentration, after which samples were diluted to 1 mg/mL in 5% SDS, 50 mM TEAB. 25 µg of protein from each sample was reduced with dithiothreitol to 2 mM, followed by incubation at 55°C for 60 minutes. Iodoacetamide was added to 10 mM and incubated in the dark at room temperature for 30 minutes to alkylate the proteins. Subsequently, phosphoric acid was introduced to achieve a concentration of 1.2%, followed by the addition of six volumes of 90% methanol containing 100 mM TEAB. Afterwards, the solution was added to S-Trap micros (Protifi) and centrifuged at 4,000 x g for 1 minute. The S-Traps containing trapped protein were washed twice by centrifuging through 90% methanol, 100 mM TEAB. 1 µg of trypsin was reconstituted in 20 µL of 100 mM TEAB and then introduced into the S-Trap, followed by the addition of 20 µL of TEAB to prevent the sample from drying out. To prevent the solution from being expelled during digestion, the cap of the S-Trap was gently secured but not fully tightened. Samples were placed in a humidity chamber at 37°C overnight. The following day, the S-Trap was centrifuged at 4,000 x g for 1 minute to collect the digested peptides. Successive additions of 0.1% TFA in acetonitrile and 0.1% TFA in 50% acetonitrile were introduced into the S-trap, followed by centrifugation and pooling. The samples were then frozen and dried using a Speed Vac (Labconco) and reconstituted in 0.1% trifluoroacetic acid before analysis.

### Mass Spectrometry

The peptides were loaded onto a 75μm × 2 cm trap column (Thermo Fisher) before being re- focused onto an Aurora Elite 75μm × 15 cm C18 column (IonOpticks) via a Vanquish Neo UHPLC system (Thermo Fisher), which was linked to an Orbitrap Astral mass spectrometer (Thermo Fisher). The composition of solvent A consisted of 0.1% formic acid in water, while solvent B comprised 0.1% formic acid in 80% acetonitrile. An Easy-Spray source operating at 2 kV was used to introduce ions to the mass spectrometer. The gradient started at 1% B and gradually rose to 5% B within 0.1 minutes, then further increased to 30% B over 12.1 minutes. Following this, it increased to 40% B within 0.7 minutes, and ultimately reached 99% B within 0.1 minutes, maintaining this composition for 2 minutes to cleanse the column, resulting in a total runtime of 15 minutes. Following the completion of each run, the column was re-equilibrated with 1% B before the next injection. The Orbitrap Astral operated in data-independent acquisition (DIA) mode, capturing MS1 scans in the Orbitrap at a resolution of 240,000, with a maximum injection time of 5 ms across a range of 380-980 m/z. An astral mass analyzer was used to generate DIA MS2 scans with a maximum injection time of 3 ms, employing a variable windowing scheme. This included 2 Da windows spanning from 380 to 680 m/z, 4 Da windows from 680 to 800 m/z, and 8 Da windows from 800 to 980 m/z. The HCD collision energy was adjusted to 25%, while the normalized AGC was configured to 500%. Fragment ions were gathered across a scan range spanning from 150 to 2000 m/z, with a cycle time of 0.6 seconds.

### Data Analysis

After sample processing and data collection, DIA-NN version 1.8.1 was used to process raw data (https://github.com/vdemichev/DIA-NN) ^19^. In all experiments, the library free analysis mode within DIA-NN was used to perform data analysis. To create annotations for the library, the mouse UniProt database, which follows the ’one protein sequence per gene’ format (UP000000589_10090), was utilized, with ’deep learning-based spectra and retention time prediction’ activated. To generate precursor ions, the settings included a maximum of 1 missed cleavage, a maximum of 1 variable modification for Ox(M), peptide lengths spanning 7 to 30, precursor charges spanning from 2 to 4, precursor m/z ranging from 380 to 980, and fragment m/z spanning 150 to 2000. Quantification was configured in ’Robust LC (high precision)’ mode with normalization set to RT-dependent, MBR enabled, protein inferences set to ’Genes’, and ’Heuristic protein inference’ disabled. The software automatically determined the MS1 and MS2 mass tolerances, as well as the scan window size. Following that, precursors were filtered based on library precursor q-value (1%), library protein group q-value (1%), and posterior error probability (50%). Protein quantification utilized the MaxLFQ algorithm within the DIA-NN R package (https://github.com/vdemichev/diann-rpackage), while the DiannReportGenerator Package (https://github.com/kswovick/DIANN-Report-Generator) was employed to count the number of peptides quantified in each protein group ^20^.

To determine proteins exhibiting significant differences between two experimental groups, a volcano plot was generated, depicting the log2 fold change (FC) against the -Log10 of the p-value. Cut-off criteria, requiring a log2FC > 1 and a -log10(p-value) > 1.3 were applied. Proteins showing significant decrease between the two groups were inputted into the PANTHER classification system ^21^ (http://pantherdb.org/) to categorize their protein types. The gene ontology classification tool DAVID)^22^ (https://david.ncifcrf.gov/) was used to determine the alterations in biological processes (BPs), molecular functions (MFs), and cellular components (CCs) associated with each condition. To understand how the significantly downregulated and upregulated proteins were interacting with each other, we analyzed the protein network using the Search Tool for Retrieval of Interacting Genes/Proteins (STRING), version 11.0 ^23^.

### Quantification of biomechanical properties of uninjured tendons

To assess the biomechanical properties of wildtype and *Cochlin*^-/-^ tendons (n=7-8 per genotype per age), tendons were harvested from the hind paws. Specifically, each flexor digitorum longus (FDL) tendon was meticulously dissected at the myotendinous junction using a dissecting microscope. Subsequently, the tarsal tunnel was incised, allowing for the gradual release and isolation of the FDL tendon. Using a dissecting microscope, any extra connective tissues, such as muscle, were carefully removed, and the flexor tendon was prepared for uniaxial testing. Two segments of 3M coarse 60-grit sandpaper were affixed to custom 3D printed grips with cyanoacrylate adhesive (Superglue, LOCTITE). Holes for steel screws were threaded using a T-handle tap wrench. Isolated tendons were placed onto grips using a custom 3D printed alignment tool, ensuring a consistent, average gauge length of 4.9 ± 0.28 mm. Tendons were bonded to the grips using cyanoacrylate adhesive and fastened with steel screws before curing overnight at in PBS at 4^°^C. All steps were conducted with the tissue intermittently immersed in PBS to prevent tissue desiccation. The cross-sectional area of the tendons was measured in Image J from digital images taken just prior to testing. The tendon’s cross-section was approximated as a rectangle, with the tissue’s width representing the major axis and the thickness representing the minor axis. Subsequently, each gripped tendon was relocated into a semi-customized uniaxial micro tester (eXpert 4000 Micro Tester, ADMET, Inc, Norwood Ma). A uniaxial displacement-controlled stretching at a rate of 0.1 mm/s until failure was implemented. Data on load, end-to-end displacement, and stress-strain were recorded and analyzed, while the failure mode was documented for each mechanically tested sample. The load-displacement and stress-strain data were graphed and used to determine both structural (stiffness) and material (modulus) properties. Specifically, the stiffness of the tested sample was determined by assessing the slope of the linear segment from the load displacement graph. The slope of the linear portion observed in the stress-strain graph was utilized as the elastic modulus parameter for each tested tendon.

### Flexor tendon injury model

Wildtype and *Cochlin^-/-^* mice (n= 8 per genotype) underwent complete transection and repair of the flexor digitorum longus (FDL) tendon as previously reported ^24–28^. Briefly, mice were administered a preoperative subcutaneous injection of 15-20 μg of sustained-release buprenorphine. Ketamine (100 mg/kg) and Xylazine (10 mg/kg) were used to induce anesthesia.

After removal of hair and sterilization of the surgical site, the FDL tendon was fully transected at the myotendinous junction (MTJ) to transiently minimize strain on the healing tendon. The FDL in the hind paw was then exposed by making a shallow incision on the posterolateral surface of the skin, and subsequently, the FDL tendon was identified and fully transected. The tendon was then repaired using 8-0 suture with a modified Kessler pattern, and the skin was closed with 5-0 suture.

### Biomechanical analysis of injured tendons

On day 28 post-surgery, animals were euthanized, and hindlimbs were harvested at the knee joint. The medial aspect of the hindlimb underwent careful dissection to free the flexor digitorum longus (FDL) tendon from both the tarsal tunnel and myotendinous junction, ensuring preservation of the repaired tendon stubs with the suture intact. The proximal end of the tendon was subsequently affixed between two sections of tape using cyanoacrylate. To assess the flexion function of the metatarsophalangeal (MTP) joint, weights ranging from 0 to 19 g were applied to the distal tendon. The flexion angle of the MTP joint was measured for each applied weight, while gliding resistance was calculated across the range of applied loads as previously described^27, 29, 30^. Reduced MTP Flexion Angle and increased Gliding Resistance indicate compromised gliding function, correlating with elevated scar tissue and peritendinous adhesion formation ^28^. After conducting the gliding tests, the FDL was freed from the tarsal tunnel. Next, the proximal end of the tendon and the digits were securely fastened within custom grips positioned opposite each other on an Instron 8841 uniaxial testing system (Instron Corporation, Norwood, MA). The tendon underwent a loading procedure at a consistent rate of 30 mm per minute until it reached failure point. The load-displacement and stress-strain data obtained were analyzed to ascertain structural properties, specifically stiffness and maximum load at failure. Stiffness of the tested sample was derived from the slope of the linear segment in the load-displacement curve.

### Statistics

Data are expressed as mean ± standard deviation with sample sizes indicated in figure legends. Unpaired Student’s t-tests with Welch’s correction, or two-way analysis of variance (ANOVA) with Šidák’s post-hoc test were used to compare differences between groups, as indicated in figure legends. A p-value ≤ 0.05 was used to define statistical significance. Statistical analyses were performed in GraphPad Prism 10 Software.

## Results

### Cochlin^-/-^ alters collagen fibril diameter distribution

To investigate whether loss of *Cochlin* affects tendon homeostasis, we examined changes in collagen fibril diameter and density using TEM. Morphologically, there was a consistent increase in collagen fibril diameter in *Cochlin^-/-^,* relative to WT tendons at all ages (**Fig. 1**). More specifically, collagen fibril diameter distribution was substantially altered between WT (median = 131.1, Q1 = 106.7, Q3 = 187.3) and *Cochlin*^-/-^ tendons (median = 170.0, Q1 = 124,.4 Q3 = 231.3), resulting in a 23% increase in collagen fibril diameter of 3-month-old *Cochlin*^-/-^ tendons compared to WT (p<0.0001) (**Fig. 1A-C**). No significant difference in collagen fibril density was observed between WT and *Cochlin*^-/-^ at 3 months (p=0.88) (**Fig. 1A & D)**.

**Figure 1.**
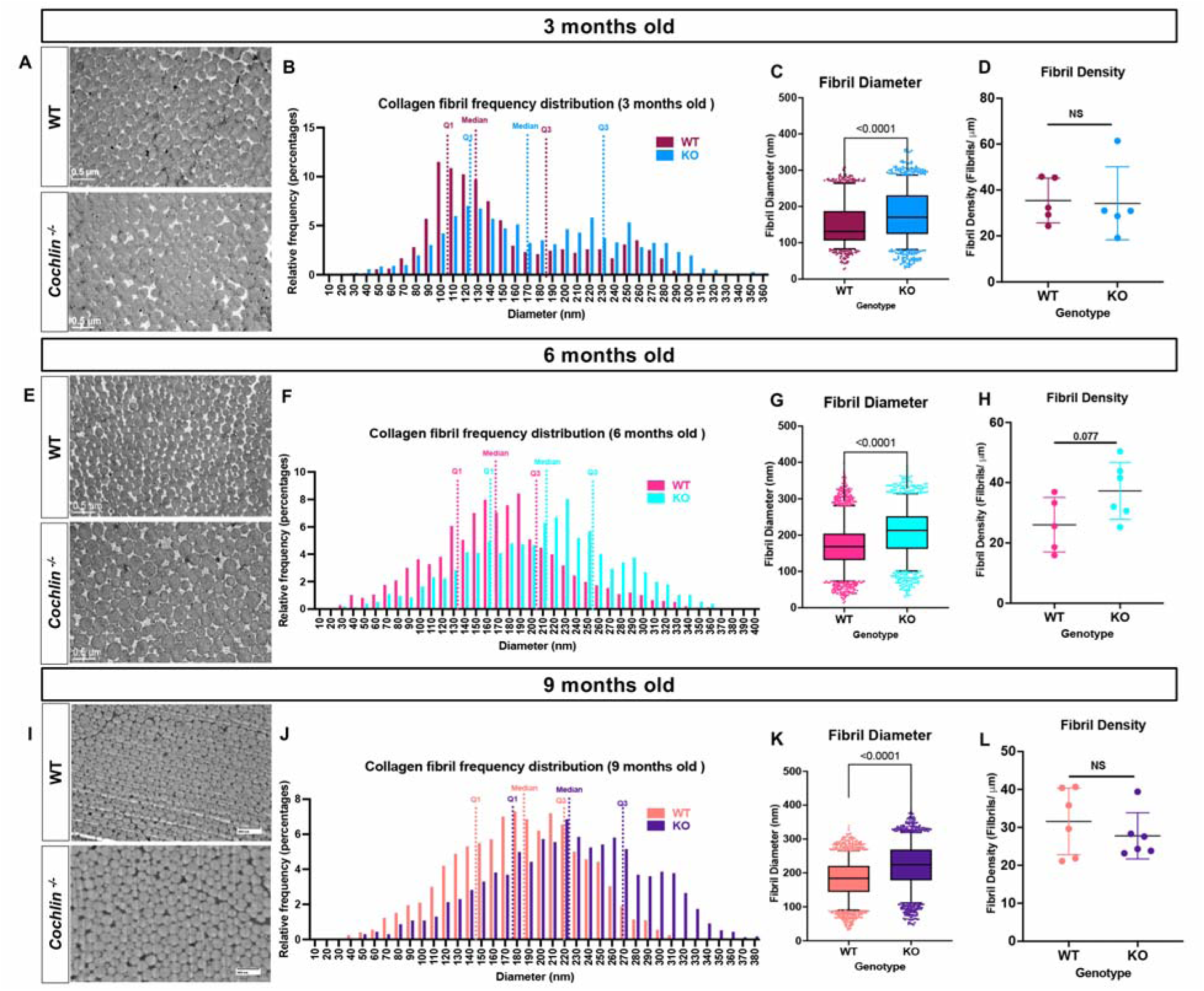
Loss of *Cochlin* drives impairments in tissue structure throughout adulthood. (**A**) Representative TEM images of FDL tendons from 3-month-old WT and *Cochlin*^-/-^ mice. (**B**) Collagen fibril diameter frequency distribution of WT and *Cochlin*^-/-^ tendons at 3 months. (**C**) Collagen fibril diameter graph shows an increase in fibril diameter of *Cochlin*^-/-^ tendons relative to WT at 3 months. (**D**) Collagen fibril density of WT and *Cochlin*^-/-^ tendons at 3 months. **(E)** Representative TEM images of uninjured WT and *Cochlin*^-/-^ FDL tendons at 6 months. (**F**) Collagen fibril diameter frequency distribution of WT and *Cochlin*^-/-^ tendons at 6 months. (**G**) Collagen fibril diameter graph shows an increase in fibril diameter of *Cochlin*^-/-^ tendons relative to WT at 6 months. (**H**) Collagen fibril density of WT and *Cochlin*^-/-^ tendons at 3 months. (**I**) Representative TEM images of uninjured WT and *Cochlin*^-/-^ FDL tendons at 9 months. (**J**) Collagen fibril diameter frequency distribution of WT and *Cochlin*^-/-^ tendons at 9 months. (**K**) Collagen fibril diameter graph shows an increase in fibril diameter of *Cochlin*^-/-^ tendons relative to WT at 9 months. (**L**) Collagen fibril density of WT and *Cochlin*^-/-^ tendons at 3 months. N= 5-6. Students Unpaired t-test with Welch’s correction used to assess statistical significance between genotypes in fibril density, and an unpaired t-test with Kolmogorov-Smirnov test was used to determine significance between genotypes in collagen fibril diameter. p < 0.05 indicates statistical significance.

At 6 months, differences in collagen fibril diameter distribution were retained between WT (median = 168.1, Q1 = 131.1, Q3 = 204.4) and *Cochlin*^-/-^ tendons (median = 129.7, Q1 = 161.9, Q3 = 252.0), with a 21% increase in fibril diameter in *Cochlin*^-/-^ relative to WT (p<0.0001) (**Fig. 1E-G**), and no differences in collagen fibril density were observed between WT and *Cochlin*^-/-^ tendons at 6 months (p=0.077) (**Fig. 1E & H**).

Finally, at 9 months, these changes in collagen fibril structure persisted. Substantial differences in fibril diameter distribution were observed between WT (median = 184.1, Q1 = 144.1, Q3 = 220.7) and *Cochlin*^-/-^ (median = 224.2, Q1 = 178.2, Q3 = 269.0) tendons, with an 18% increase in the median collagen fibril diameter in *Cochlin*^-/-^ tendons relative to WT (p<0.0001) (**Fig. 1I-K**). As with younger ages, no significant difference in collagen fibril density were observed between genotypes at 9 months (p=0.45) (**Fig. 1I & L**).

### Cochlin^-/-^ alters tendon biomechanical properties at 6 and 9 months

Based on the progressive changes in collagen matrix structure with *Cochlin^-/-^,* we then assessed tendon material and structural mechanical properties at 6 and 9 months. No significant differences in cross-sectional area (CSA) or elastic modulus were observed between *Cochlin*^-/-^ and WT tendons at either 6-months (CSA: p=0.81), elastic modulus: p= 0.32) or 9-months (CSA: p= 0.42, elastic modulus: p= 0.29) of age (**Fig. 2A & B**). In contrast, significant decreases in tendon stiffness were observed in *Cochlin*^-/-^ tendons, relative to WT at 6 months (WT: 7.43 ± 3.236 N/mm, KO: 5.51 ± 2.528 N/mm, p=0.0064), although no significant difference was observed at 9 months (WT: 7.599 ± 1.226 N/mm, KO: 6.15 ± 1.051 N/mm, p=0.0724) (**Fig. 2C**). Moreover, *Cochlin*^-/-^ displayed significant decreases in peak load at both 6 months (WT: 10.09 ± 3.342 N, KO: 7.64 ± 2.802 N, p=0.0171) and 9 months (WT: 10.52 ± 2.874 N, KO: 8.07 ± 4.824 N, p=0.0042) (**Fig. 2D**) compared to WT, further supporting disruptions in tendon homeostasis with *Cochlin*^-/-.^

**Figure 2.**
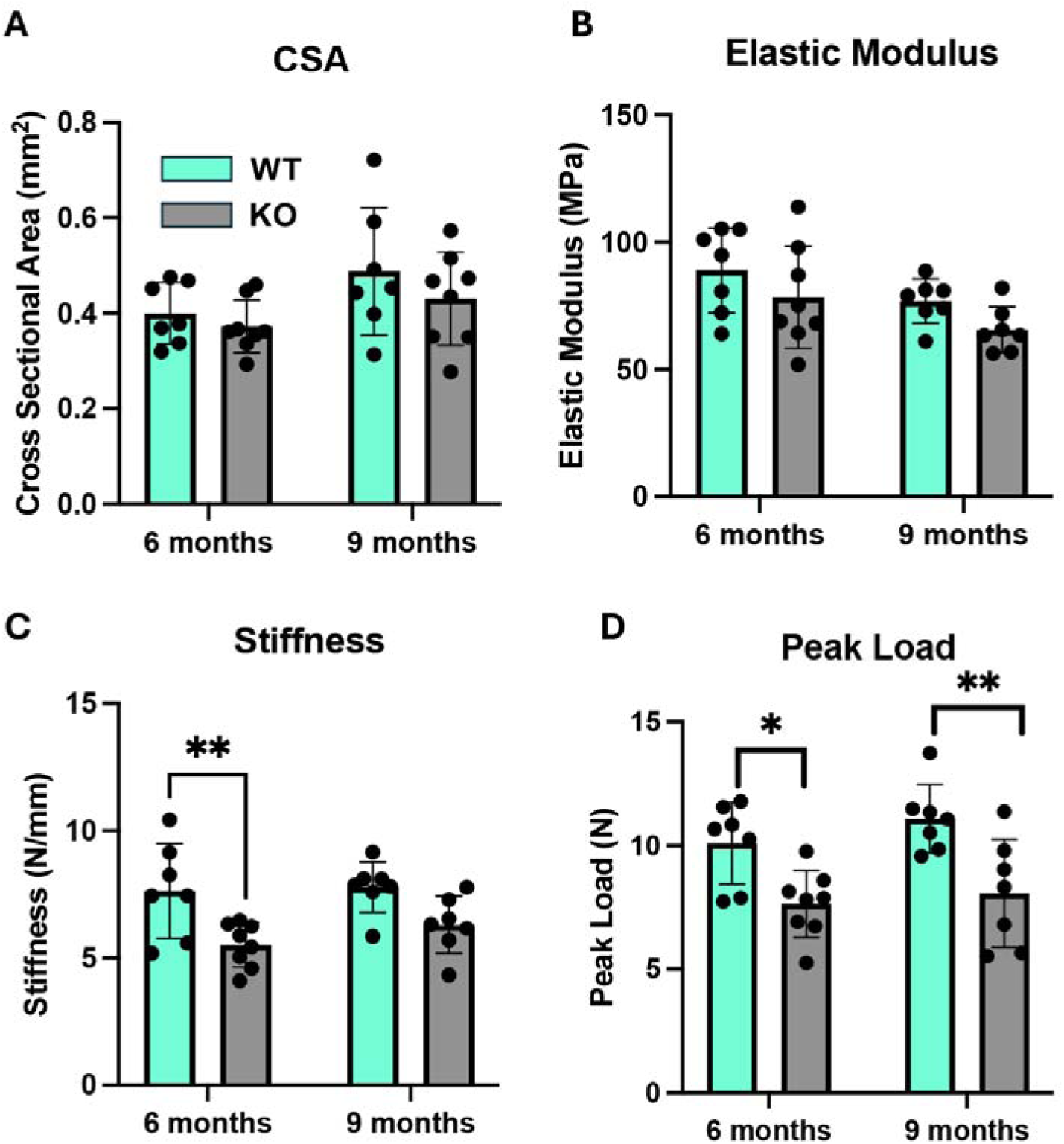
Loss of *Cochlin* results in structural impairments. Measurement of (**A**.) Cross-sectional area (CSA), (**B**) Elastic Modulus, (**C**) Stiffness, and (**D**) Peak Load of 6- and 9-month-old WT and *Cochlin*^-/-^ flexor tendons. N= 7-8. Two-way ANOVA with Šidák’s multiple comparison test was used to assess statistical significance between genotypes. (*) indicates p<0.05. (**) p<0.01.

### Loss of Cochlin drives differential expression of several proteins at 3 and 6 months

To better understand both how *Cochlin*^-/-^ alters of the composition of the tendon proteome, and to begin to better understand the mechanisms underpinning the dysregulated collagen structure and impaired mechanics in *Cochlin*^-/-^ tendons, we conducted proteomic analyses of WT and *Cochlin*^-/-^ tendons at 3 months and 6 months of age. In 3-month-old tendons, 6 proteins were significantly downregulated in *Cochlin*^-/-^ relative to WT, while 21 proteins were significantly upregulated in *Cochlin*^-/-^ tendons relative to WT (**Fig 3**. A & B). The proteins that were downregulated in *Cochlin*^-/-^ tendons were associated with RNA metabolism (32%), ECM proteins (20%), cytoskeletal proteins (28%), and transmembrane signal receptor proteins (20%) (**Fig. 3 C**). The biological processes associated with these downregulated genes were *Sensory perception of sound* (GO: 0007605) and *Positive regulation of neuron projection development* (GO: 0010976) (**Fig. 3D**). Differentially expressed proteins were then analyzed using the STRING database to construct a protein interaction network. We did not observe interactions between any of the downregulated proteins in 3-month-old *Cochlin*^-/-^ tendons (**Fig. 3E**). In terms of the 21 proteins upregulated in *Cochlin*^-/-^ tendons, 40% were classified as protein modifying enzymes, 20% as metabolite interconversion enzymes, 20% protein-binding activity modulators, and 20% as response to stimulus proteins (**Fig. 3F**). The biological processes associated with proteins upregulated in *Cochlin*^-/-^ tendons include *positive regulation of cell proliferation* (GO: 0008284), *positive regulation of fat cell differentiation* (GO: 0045600), and *lysosome organization* (GO: 0007040) (**Fig. 3G**). When we looked at the interaction between these upregulated proteins, we observed one notable interaction involving Rnaseh2b and RFC3 (**Fig. 3H**). Rnaseh2b is a subunit of the ribonuclease H2 enzyme complex, and RFC3 is a subunit of the replication factor C (RFC) complex, which is essential for DNA replication and repair ^31^.

**Figure 3.**
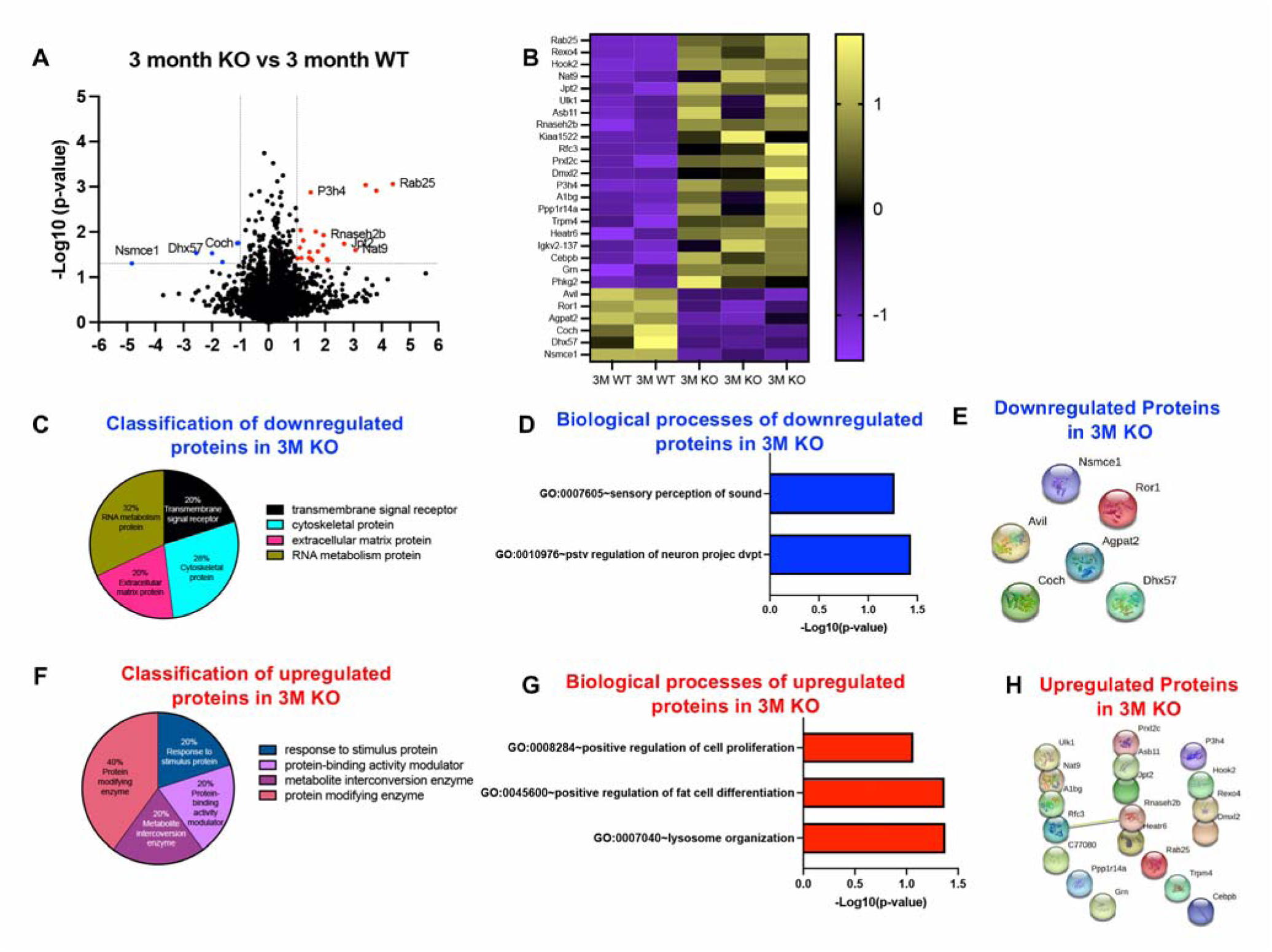
Loss of *Cochlin* results in impairments in tendon homeostasis at 3 months. (**A**) Volcano plot and (**B**) Heatmap showing the significantly different protein abundances between WT and *Cochlin*^-/-^ tendons at 3 months. (**C**) Classification of all downregulated proteins in *Cochlin*^-/-^ tendons relative to WT tendons at 3 months. (**D**) Biological process downregulated in *Cochlin*^-/-^ tendons relative to WT tendons. (**E**) Protein-protein interaction of all the downregulated proteins between WT and *Cochlin*^-/-^ groups at 3 months. (**F**) Classification of all upregulated proteins in *Cochlin*^-/-^ tendons relative to WT tendons at 3 months. (**G**) Biological processes upregulated in *Cochlin*^-/-^ tendons relative to WT tendons. (**H**) Protein-protein interaction of all the upregulated proteins between WT and *Cochlin*^-/-^ groups at 3 months.

In 6-month-old tendons we identified 14 proteins that were significantly downregulated in *Cochlin*^-/-^ tendons and 48 proteins that were significantly upregulated in *Cochlin*^-/-^ tendons relative to WT (**Fig. 4A & B**). Classification of all downregulated proteins revealed that 40% were RNA metabolism proteins, 20% were ECM proteins, 20% were membrane traffic proteins, and 20% were cytoskeletal proteins (**Fig. 4C**). Biological processes associated with these downregulated proteins included *sensory perception of sound* (GO: 0007605), *rRNA processing* (GO: 0006364), *ribosome biogenesis* (GO: 0042254), and *sarcomere organization* (GO:0045214) (**Fig. 4D**). STRING analysis identified two different groups of interaction in the downregulated proteins set, including interactions between RFP2 and DCAF13 (**Fig. 4E**). While the functional significance of this interaction is unknown, RFP2 is involved in ER-associated degradation processes^32^, and DCAF13 is involved in rRNA processing^33^. The proteins that were upregulated in *Cochlin*^-/-^ tendons were classified as metabolite interconversion enzymes (26.2%), Protein binding activity modulators (10.5%), Protein modifying enzymes (10.4%), Intercellular signal molecules (10%), membrane traffic proteins (5.3%), cell adhesion molecules (5.3%), transfer/carrier proteins (5.3%), transmembrane signal receptors (5.3%), response to stimulus proteins (5.3%), cytoskeletal proteins (5.2%), scaffold/adaptor proteins (5%), and transporter proteins (5%) (**Fig. 4F**). Biological processes associated with the proteins upregulated in

**Figure 4.**
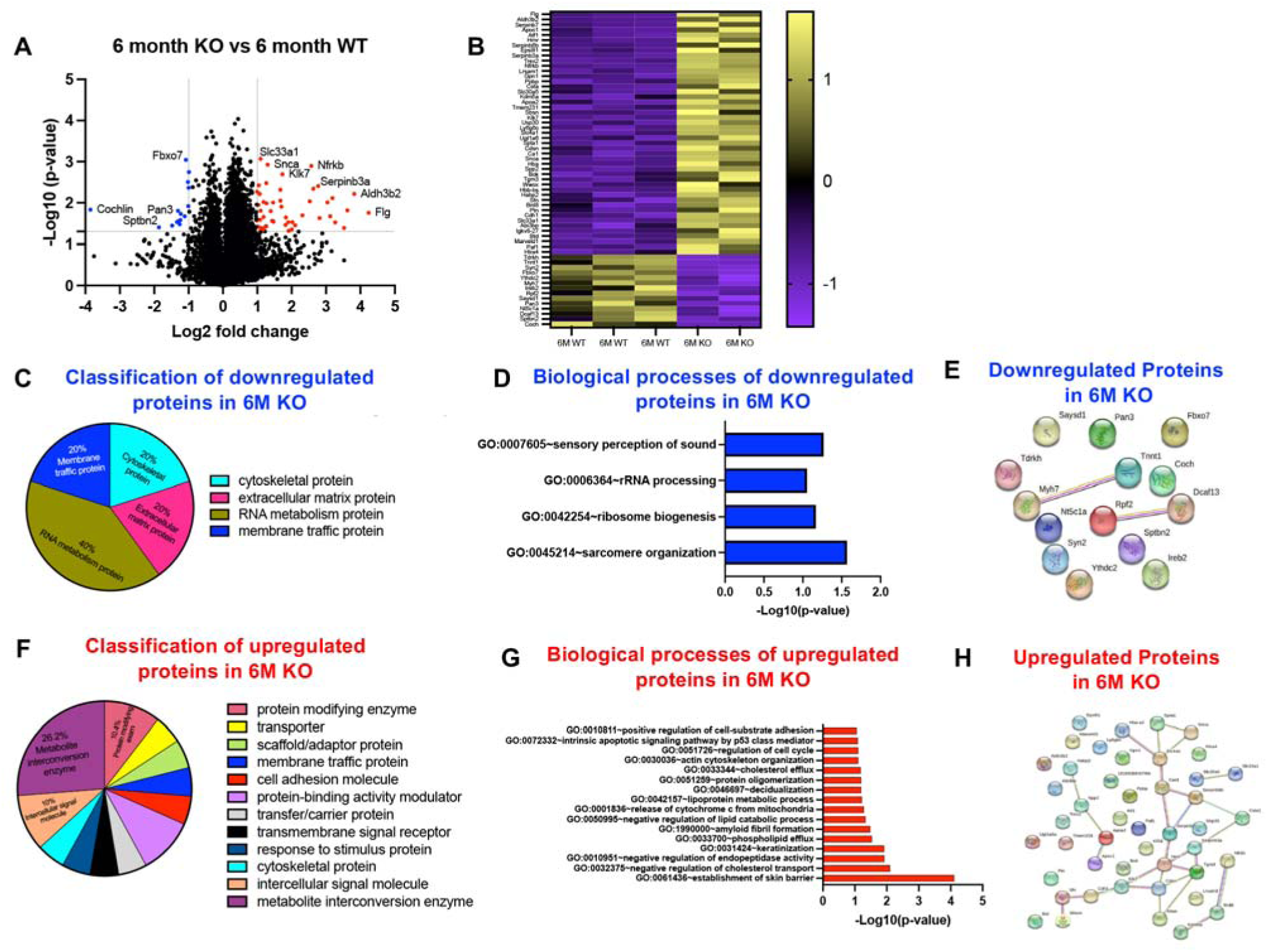
Loss of *Cochlin* results in impairments in tendon homeostasis at 6 months. (**A**) Volcano plot and (**B**) Heatmap showing the significantly different protein abundances between WT and *Cochlin*^-/-^ tendons at 6 months. (**C**) Classification of all downregulated proteins in *Cochlin*^-/-^ tendons relative to WT tendons at 6 months. (**D**) Biological process downregulated in *Cochlin*^-/-^ tendons relative to WT tendons. (**E**) Protein-protein interaction of all the downregulated proteins between WT and *Cochlin*^-/-^ groups at 6 months. (**F**) Classification of all upregulated proteins in *Cochlin*^-/-^ tendons relative to WT tendons at 6 months. (**G**) Biological processes upregulated in *Cochlin*^-/-^ tendons relative to WT tendons. (**H**) Protein-protein interaction of all the upregulated proteins between WT and *Cochlin*^-/-^ groups at 6 months.

### Cochlin^-/-^ tendons included protein oligomerization (GO: 0051259), actin cytoskeleton

*organization* (GO: 0030036), and *positive regulation of cell-substrate adhesion* (GO: 0010811) (**Fig. 4G**). Furthermore, STRING analysis identified 4 different groups of interactions in the proteins upregulated in *Cochlin*^-/-^ tendons at 6 months of age (**Fig. 5H**). The first interaction set included APOC1, APOA2, SPP2, and HABP2, which are involved in extracellular matrix organization and lipid metabolism. The second set of interaction was observed between KDM5A, BRD8, and NFRKB which are related to chromatin remodeling and transcriptional regulation. The third interaction was observed between SLC30A5 and SLC33A1 which are involved in metal ion transport and cellular homeostasis. Finally, the fourth set of interactions was observed between 14 proteins including WWOX, which plays a direct role in cellular apoptosis and TGFB1 mediated cell death, and SFN which is involved in cell cycle regulation. The extensive interaction network among these 14 proteins highlights their diverse roles in cellular structure, function, and homeostasis and reflects a complex response to loss of *Cochlin*, potentially impacting processes such as apoptosis, cell signaling, cell adhesion, chromatin remodeling, proteolysis, cytoskeletal integrity, and ECM organization.

**Figure 5.**
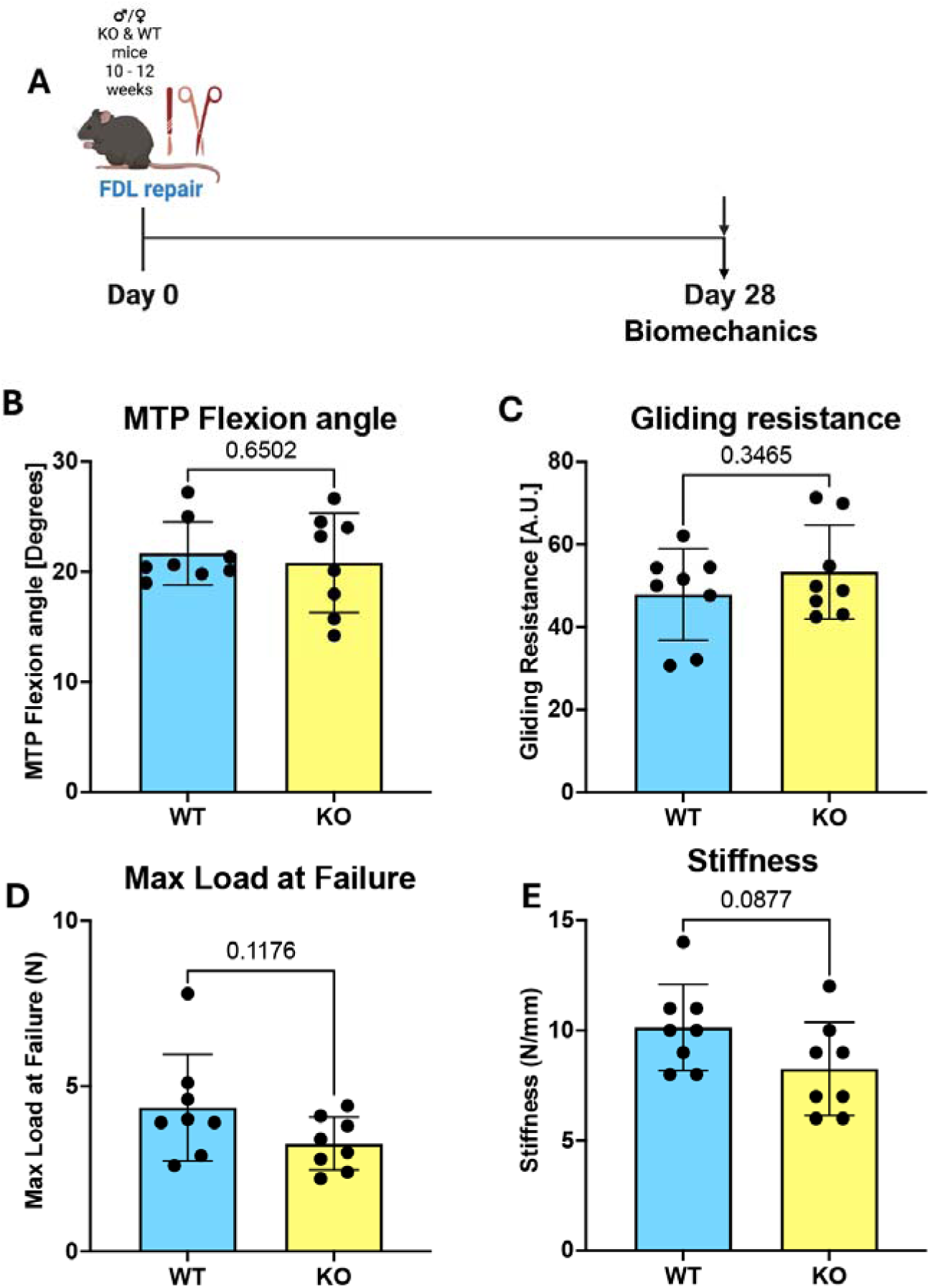
Loss of *Cochlin* does not affect tendon healing. (**A**) Schematic of experimental timeline. Measurement of (**B**) metatarsophalangeal (MTP) joint flexion angle, (**C**) Gliding resistance, (**D**) Maximum load at failure and (**E**) Stiffness of WT and *Cochlin*^-/-^ flexor tendons isolated after 28 days of flexor tendon injury and repair. N = 7-8. An unpaired t-test with Welch’s correction used to assess statistical significance between genotypes.

### Loss of Cochlin does not impact tendon healing

Finally, we aimed to determine the effects of *Cochlin*^-/-^ on the tendon healing process. We performed flexor tendon transection and surgical repair on 10–12-week-old WT and *Cochlin*^-/-^ mice and assessed mechanical and functional outcomes after 28 days (**Fig. 5A**). Despite alterations in ECM structure at 3 months, the absence of C*ochlin* did not impair the healing process, as no changes in MTP flexion angle (p = 0.6502; WT vs. *Cochlin*^-/-^) or gliding resistance (p =0.3465; WT vs *Cochlin*^-/-^) were observed (**Fig. 5B & C**). Additionally, there were no differences in maximum load at failure (p=0.117) or stiffness (p=0.08) between genotypes (**Fig. 5D & E**). These findings suggest that *Cochlin* is required for maintaining tendon homeostasis but not for the healing process.

## Discussion

Tendon homeostasis is crucial for preserving tissue integrity and function, enabling tendons to resist mechanical strain and maintain optimal performance over time ^2, 4, 5^. Disruptions in this balance, whether due to mechanical overload, aging, or disease, can lead to tendon degeneration and increased susceptibility to injury ^2, 4^. Furthermore, compromised tendon homeostasis is associated with impaired collagen structure and organization, and altered cell activity, ultimately weakening the tendon’s mechanical properties ^1, 5, 9^. Given that a consistent loss of Cochlin protein and *Cochlin*+ cells was observed in two different models of disrupted homeostasis (natural aging and depletion of Scleraxis-lineage cells in young adult mice)^4^, we defined the requirement for Cochlin in maintaining tendon homeostasis in terms of collagen fibril structure, and tissue mechanics, and determined how loss of Cochlin alters the tendon proteome.

Cochlin plays a critical role in maintaining the structural integrity and function of several tissues ^12, 13, 17^. Although the mechanisms through which Cochlin maintains tissue structure are unclear, Cochlin has multiple functional domains ^34^, including two von Willebrand factor A-like (vWFA) domains, which have high-affinity for collagen ^35^, and thus may facilitate structural stability. Conversely, aberrant expression of Cochlin is associated with pathologies involving altered ECM fluid flow and shear stress, including Glaucoma ^11, 36^, suggesting the importance of maintaining Cochlin expression at physiological tissue-specific levels to support tissue homeostasis. Related to this, there has been very little prior work to define the role of Cochlin in the tendon despite Cochlin being one of the most differentially expressed genes in healthy flexor tendon relative to healing^24^ or degenerative ^37^ tendons.

To assess the role of *Cochlin* in maintaining tendon homeostasis, we first looked at how tendon structure and function were impacted after loss of *Cochlin*. The results of this study demonstrated that collagen fibril formation in tendons was significantly disrupted in the absence of *Cochlin*. In *Cochlin*^-/-^ tendons, there was a consistent skewing toward larger diameter fibrils compared to WT. Based on concomitant impairments in mechanical properties, these data suggest that the shift toward larger diameter fibrils is indicative of disrupted tissue homeostasis and structure. Moreover, given the tightly regulated process of collagen fibrillogenesis^38–44^ and the consistent loss of Cochlin from embryonic development and tendon patterning in *Cochlin*^-/-^ mice, these data suggest a requirement for Cochlin in tendon collagen fibrillogenesis.

Dysregulation in any step of fibrillogenesis can result in abnormal fibril morphology, leading to compromised tendon mechanical properties. While the precise mechanisms through loss of Cochlin alters collagen fibril structure, comparative proteomic analysis identified potential regulators of this phenotype. For example, in 3-month-old *Cochlin*^-/-^ tendons decreased expression of cytoskeletal proteins are observed which can disrupt actin bundling ^45^, subsequently affecting cell shape, motility, and structural integrity ^45^. Moreover, downregulation of catalytic enzymes including kinases, which are known to disrupt signal transduction pathways, thus affecting processes like cell cycle regulation and growth, differentiation, and metabolism ^46–48^. Collectively, these data suggest that loss of Cochlin may alter how tendon resident cells sense and interact with the extracellular matrix environment, resulting in altered cell behavior. However, it is not known whether loss of Cochlin directly alters cell function to drive changes in fibril structure, or if changes in cell function are secondary to the altered composition of the ECM. Related to this, tendons from 6-month-old *Cochlin*^-/-^ mice demonstrate upregulation of proteins related to protein binding activity, intercellular signal molecules, membrane traffic proteins, and cell adhesion molecules, suggesting a concerted cellular response to compensate for altered collagen fibril morphology and impaired mechanical function. For example, we observed an increase in cadherins, and other cell adhesion molecules involved in calcium-dependent cell adhesion^49, 50^ suggesting enhanced cell-cell adhesion. Cadherins play a role in maintaining the structure of tissues where collagen fibrils provide mechanical support ^51–54^. Consistent with an increase in cell-adhesion proteins was an increase in membrane traffic proteins and transfer/carrier proteins which may enhance the secretion of collagen precursors and other ECM components, as well as support the delivery of necessary substrates and cofactors for collagen synthesis and cross-linking respectively ^55^, consistent with an increase in collagen fibril diameter in *Cochlin*^-/-^. Moreover, increased activity of protein modifying enzymes may be linked to ECM remodeling, as modifying enzymes can activate or deactivate other proteins involved in collagen synthesis and fibril organization^56^.

Despite defining a requirement for *Cochlin* in tendon homeostasis and collagen organization, loss of Cochlin did not alter the tendon healing process. Consistent with this, we have previously demonstrated that while robust in healthy tendons, there is minimal *Cochlin* expression in the context of tendon healing ^4, 24^. These data are in line with our prior work that suggests that underlying disruption in tendon homeostasis do not necessarily lead to tendon healing deficits. For example, depletion of Scleraxis-lineage cells in young mice drives a phenotype consistent with acceleration of natural aging ^4^. However, healing in these mice is improved relative to WT littermates ^2, 4^, suggesting a disconnect between some aspects of tendon homeostasis and the response to injury.

In terms of limitations of this study, mice with global and non-inducible deletion of Cochlin were used. While this provides an opportunity to define the role of Cochlin in tendon development and subsequent post-natal growth and adult homeostasis, it does not recapitulate the likely progressive loss of Cochlin that occurs during natural aging. Moreover, our prior work has identified a consistent loss of other ECM components during disrupted homeostasis (e.g., Chondroadherin, Keratocan, and Aggrecan ^4^), and it is likely that more profound phenotypes would be observed in compound mutants. Finally, this approach does not result in tendon-specific deletion of Cochlin. While these mice do not display any overt phenotypes, increases susceptibility to bacterial infections, and decreased cytokine production in response to bacterial infection have been reported ^57, 58^. As such, subsequent work using some combination of inducible, tendon-specific conditional deletion constructs and/or combined deletion of multiple ECM components should be considered.

Taken together, this study demonstrates the requirement for Cochlin in maintaining tendon homeostasis, including development or preservation of collagen fibril organization, tendon mechanics, and maintenance of a normal tendon proteome. Moreover, this work further supports the concept that disruptions in tendon homeostasis do not necessarily lead to impairments in the tendon healing process. Collectively, these data identify Cochlin as a critical regulatory component of proper tendon structure and future work will define the therapeutic potential of conservation or restoration of Cochlin to facilitate continued tendon health through the lifespan.

## Funding

This work was supported in part by NIH/NIAMS R01 AR082667 (AEL), R00 AR080757 (AECN), P30 AR069655, and the Schlumberger Foundation Faculty for the Future Fellowship (EAS).

## Acknowledgements

We would like to acknowledge the technical support of the Electron Microscopy Resource and the Mass Spectrometry Resource in the Center for Advanced Research Technology, and the Biomechanics and Multimodal Tissue Imaging Core in the Center for Musculoskeletal Research at the University of Rochester Medical Center

## Author contributions

Conceptualization: EAS, AEL; Funding acquisition: ES, AECN, AEL; Methodology: EAS, EL, AECN, MRB; Formal analysis: EAS, EL, NA, CS, Writing-original draft: EAS, EL, AEL; Writing-review and editing: EAS, EL, NA, CS, AECN, MRB, AEL.

## Notes

### Competing Interest Statement

The authors have declared no competing interest.

